# Hormone Control Regions mediate opposing steroid receptor-dependent genome organizations

**DOI:** 10.1101/233874

**Authors:** François Le Dily, Enrique Vidal, Yasmina Cuartero, Javier Quilez, Silvina Nacht, Guillermo P. Vicent, Priyanka Sharma, Gaetano Verde, Miguel Beato

## Abstract

In breast cancer cells, topologically associating domains (TADs) behave as units of hormonal gene regulation with transcripts within hormone responsive TADs changing coordinately their expression in response to steroid hormones. Here we further described that responsive TADs contain 20-100 kb-long clusters of intermingled estrogen receptor (ER) and progesterone receptor (PR) binding sites, hereafter called Hormone-Control Regions (HCRs). We identified more than 200 HCRs, which are frequently bound by ER and PR even in the absence of hormones. These HCRs establish steady long-distance inter-TAD interactions between them and organize characteristic looping structures with promoters even in the absence of hormones. This organization is dependent on the expression of the receptors and is further dynamically modulated in response to steroid hormones. HCRs function as platforms integrating different signals resulting in some cases in opposite transcriptional responses to estrogens or progestins. Altogether, these results suggest that steroid hormone receptors act not only as hormone-regulated sequence-specific transcription factors, but also as local and global genome organizers.

**Highlights:**

Hormone responsive TADs are organized around conserved large regulatory
regions (HCRs) enriched in ER and PR.
HCR contact promoters within their TADs and engaged long-range inter-TADs
contacts between them.
Binding of the receptors in absence of hormones maintains global HCR-HCR
interactions and intra-TADs regulatory loops.
HCRs can integrate the hormone signals in divergent ways leading to opposite
restructuration of TADs in response to Estrogens or Progestins.

## Introduction

The folding of the eukaryotic chromatin fiber within the cell nucleus together with nucleosome occupancy, linker histones and post-translational modifications of histones tails, plays an important role in modulating the function of the genetic information. It is now well demonstrated that the genome is organized non-randomly in a hierarchy of structures with chromosomes occupying territories that are partitioned into segregated active and inactive chromatin compartments (Fraser et al. 2015). Furthermore, chromosomes are segmented into contiguous “topologically associating domains” (TADs), within which chromatin interactions are stronger than with the neighboring regions (Dixon et al. 2012; Nora et al. 2012). Such organization has been shown to participate in DNA replication and transcription (Pope et al. 2014; Lupianez et al. 2015). Boundaries between TADs are conserved among cell types and enriched for cohesins and CTCF binding sites as well as for highly expressed genes. However, how boundaries are established and maintained is not yet fully understood (Hou et al. 2012; Jin et al. 2013; Dixon et al. 2015). TADs are further organized in sub-domains and loops that also depend on CTCF and other factors linked to transcription regulation (Phillips-Cremins et al. 2013; Rao et al. 2014). The sub-TAD organization is more divergent between cell types and it dynamically reorganizes during the process of differentiation (Ji et al. 2016). In terminally differentiated cells, it is still not totally clear whether TADs are relatively stable pre-organized structures or if they are dynamically remodeled in response to transient external cues (Jin et al. 2013; Le Dily et al. 2014; Kuznetsova et al. 2015). In any case, it is now accepted that TADs facilitate contacts between genes promoters and their regulatory elements located away on the linear genome. However, it is not clear to what extent cell-specific transcription factors modulating the activity of those regulatory sites are also involved in organizing this particular level of chromatin folding.

Steroid receptors are stimuli induced transcription factors, which regulate the expression of thousands of genes in hormone responsive cells (Cicatiello et al. 2004; Bain et al. 2007). Notably, the Estrogen and Progesterone Receptors (ER and PR, respectively) are known to bind either directly to the promoter of their target genes or to enhancer elements where they orchestrate the recruitment of chromatin remodeling complexes and general transcription factors (Carroll et al. 2005; Hsu et al. 2010; Ballare et al. 2013; Li et al. 2013). Several studies have analyzed the effects of steroids on the 3D organization of chromatin at limited resolution leading to apparently contradictory results. For example, we previously showed that TADs can respond as units to the hormone signals with dynamic reorganization of the entire TAD (Le Dily et al. 2014). In contrast, other studies suggested that enhancers and promoters contacts precede receptor activation (Hakim et al. 2009; Jin et al. 2013). These scenarios are not mutually exclusive and it is possible that different regulatory mechanisms are required depending on the general chromatin context (Kuznetsova et al. 2015).

To gain insights into the 3D organization of chromatin in response to steroid hormones, we studied at high resolution the organization of a TAD where genes are coordinately repressed by Progestins (Pg) but activated by Estradiol (E2). We observed that within this TAD, which contains the *ESR1* gene among others, promoters were organized around a cluster of ER and PR binding sites that we term Hormone Control Region (HCR). Based on the analysis of clustered binding of ER and PR we identified 211 additional putative HCRs throughout the genome of T47D breast cancer cells. These regions had frequent interactions with promoters within their respective TAD as well as long-range HCR-HCR inter-TAD interactions. Furthermore, we observed important differences in the internal structure of HCR-containing TADs between cells expressing or not hormone receptors. Depletion of the endogenous ER and/or PR points towards a role of steroid receptors in maintaining a functional intra-TAD organization in absence of hormone. Finally, we also observed that the activity and interactions of the HCRs were dynamically modified upon exposure to steroid hormones. Notably, a subset of HCRs are acting as enhancers or silencers depending on the hormone signal received by the cells, in correlation with structural modifications of their respective TADs. Overall, these observations suggest that steroid hormone receptors act not only as hormone regulated sequence-specific transcription factors, but also as local and global genome organizers.

## Results

### The TAD encompassing the *ESR1* gene is organized around an HCR

In a previous study with T47D breast cancer cells, we observed that TADs can behave as units of response to steroid hormones (Le Dily et al. 2014). One of these steroid-responsive TADs contains the *ESR1* gene (encoding the ERα protein) and 5 other protein-coding genes, which are coordinately up-regulated by E2 and down-regulated by Pg ((Le Dily et al. 2014) and Supplementary Figure 1A). To study the organization of this domain (hereafter referred to as ESR1-TAD) at high resolution, we designed capture enrichment probes to cover a 4 Mb region of chromosome 6 including the ESR1-TAD (Figure 1A). These probes were used in Capture-C experiments to generate contact maps at various resolutions from cells grown in conditions depleted of steroid hormones (Figure 1A, 1B). We used these datasets to create virtual 4C profiles taking as baits the promoters of protein coding genes laying in ESR1-TAD (Supplementary Figure 1B). We observed that, in addition to establishing contacts between them, promoters were engaged in frequent interactions with a 90 kb intergenic region located upstream of the ESR1 gene promoter (Figure 1B, Supplementary Figure 1B). Similarly, virtual 4C profile tacking as bait the whole 90 kb intergenic region confirmed that it engaged interactions with virtually all upstream and downstream promoters of protein coding genes as well as with other non-annotated sites marked by H3K4me3 and RNA-Polymerase II (RNA Pol. II) within the boundaries of ESR1-TAD (Figure 1C). ChIP-Seq data obtained with T47D cells exposed to E2 or Pg demonstrated that the 90 kb region is enriched in binding sites for the two steroid receptors (Figure 1C, Supplementary Figure 1C). In absence of steroids (−H), ER but not PR was already bound to chromatin at 4 sites within the region (Figure 1C, Supplementary Figure 1C). Exposure of cells to E2 (+E2) led to an increase in ER binding at the pre-existing sites and only a few sites were bound *de novo* by the hormone-activated ER (Figure 1C, Supplementary Figure 1C). Upon treatment with Pg (+Pg), PR bound to approximately 40 different locations distributed throughout the ESR1-TAD (Figure 1C). Among those sites, 10 were concentrated within the 90 kb intergenic region that contacts all promoters (Figure 1C, Supplementary Figure 1C). This region is further characterized by high density of RNA Pol. II, BRD4, H3K4me3 and H3K27ac (Figure 1C) and by the expression of several non-annotated transcripts, which, similarly to the nearby protein-coding genes, were oppositely regulated by E2 and Pg ((Le Dily et al. 2014) and Supplementary Figures 1D, 1E). Upon exposure to Pg and binding of the PR, the levels of H3K27ac, BRD4 and RNA Pol II decreased within the region (Figure 1C); in correspondence with the decreased expression of the genes in this condition (Supplementary Figure 1A).

**Figure 1:**
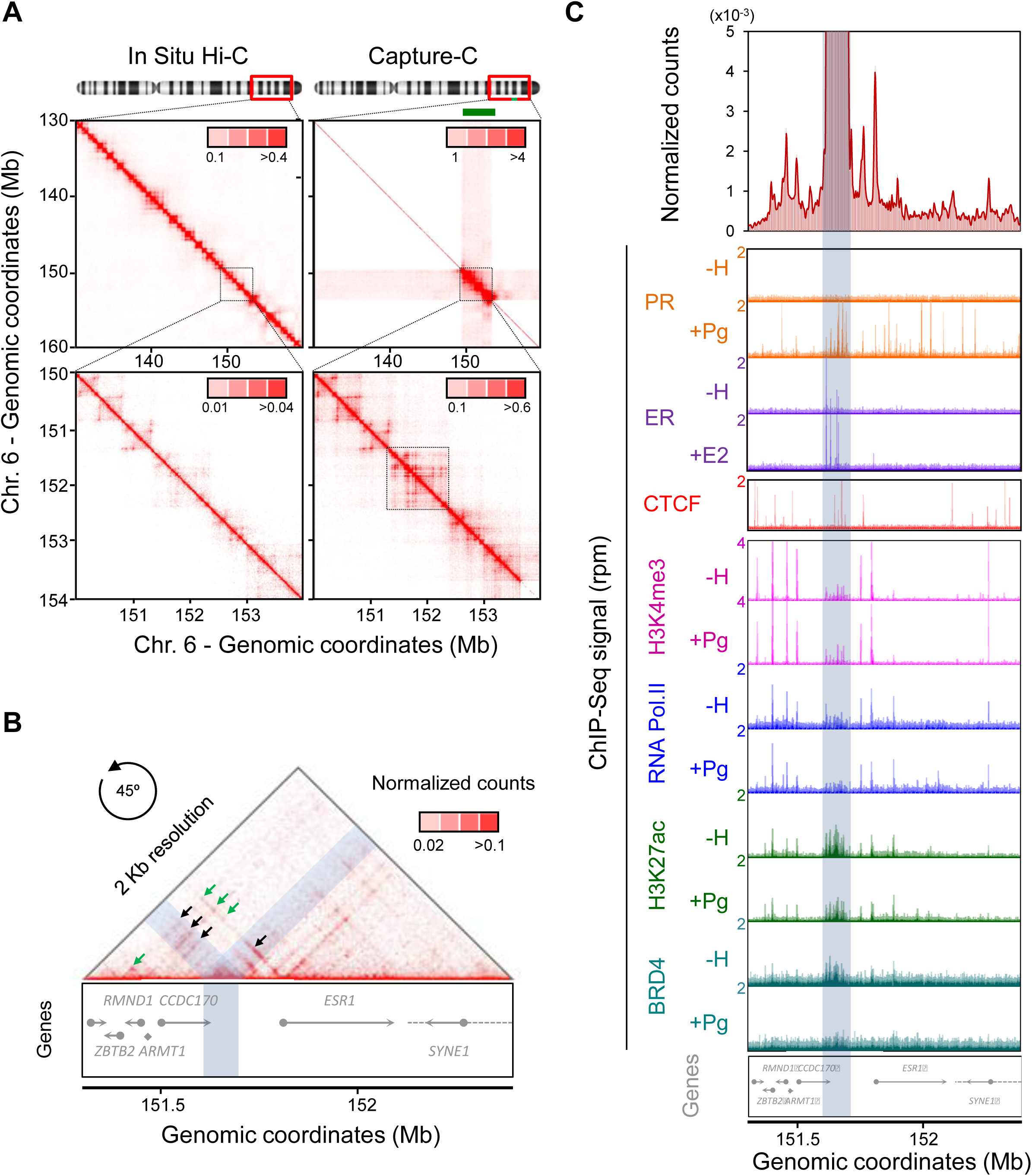
The ESR1-TAD is organized around a HCR. **(A)** *Top panels*: normalized contact maps at 100 kb resolution of a 30 Mb region of chromosome 6 encompassing the ESR1-TAD (Chr.6: 130,000,000-160,000,000) in T47D before (*in situ* Hi-C, left panel) or after capture enrichment (right panel) with biotinylated probes directed against the region marked in green (Chr.6: 149,750,000-153,750,000). *Bottom panels*: magnification at 10 kb resolution of the 4 Mb region containing the ESR1-TAD (dashed square). Color scales in insets correspond to the normalized counts number per million of genome-wide valid pairs. **(B)** Contact matrix at 2 kb resolution of the ESR1-TAD (Chr.6: 151,300,000-152,400,000; the matrix was rotated 45° and the top half is shown). Green and black arrows highlight the loops established between promoters and between promoters and a 90 kb intergenic region (highlighted in light blue), respectively. **(C)** *Top panel*: virtual 4C profile at 2 kb resolution using the 90 kb intergenic region (highlighted in blue) as bait. Panels below correspond to Genome Browser tracks of ChIP-Seq read per million (rpm) profiles obtained for the different factors and epigenetic marks listed in absence (−H) or presence of Progestin (+Pg) or Estradiol (+E2). Positions of protein coding genes and genomic coordinates are shown at the bottom.

Together, these observations indicate that, in the absence of steroids, the ESR1-TAD is organized around a 90 kb intergenic region where ER is already bound in absence of hormones and where both ER and PR cluster after exposure to their cognate ligand. We therefore designate such a region as a Hormone Control Region (HCR), which coordinates the hormone-induced changes in transcription of the genes within ESR1-TAD.

### HCRs participate in organizing the T47D genome

Next, we used ChIP-Seq datasets of ER and PR in T47D cells exposed to the cognate hormones to identify potential HCRs genome-wide. Applying the detection scheme depicted in Figure 2A (See also Methods section), we identified a total of 2681 regions enriched mainly in PR (PR≥3; ER<3), 164 enriched mainly in ER (PR<3; ER≥3) and 212 putative HCRs enriched in both ER and PR (PR≥3; ER≥3). These HCRs were located in TADs containing genes related with hormone responsive mammary cell identity, as for example *PGR* gene, which encodes the PR proteins, as well as *FOXA1* (Figure 2A). In steroid-depleted conditions these HCRs, of a median size of 30 kb, were characterized by high levels of FOXA1 binding as well as by enhancer associated marks (*e.g.,* H3K27ac, BRD4, and RNA Pol. II) as compared to their flanking regions (Figure 2B). Although HCRs were defined based on the number of binding sites after exposure to the hormones, they were frequently characterized by binding of unliganded receptors. In hormone-depleted conditions, 78% and 53% of the HCRs were already occupied at one or more site(s) by unliganted ER or PR, respectively (Figure 2C).

**Figure 2:**
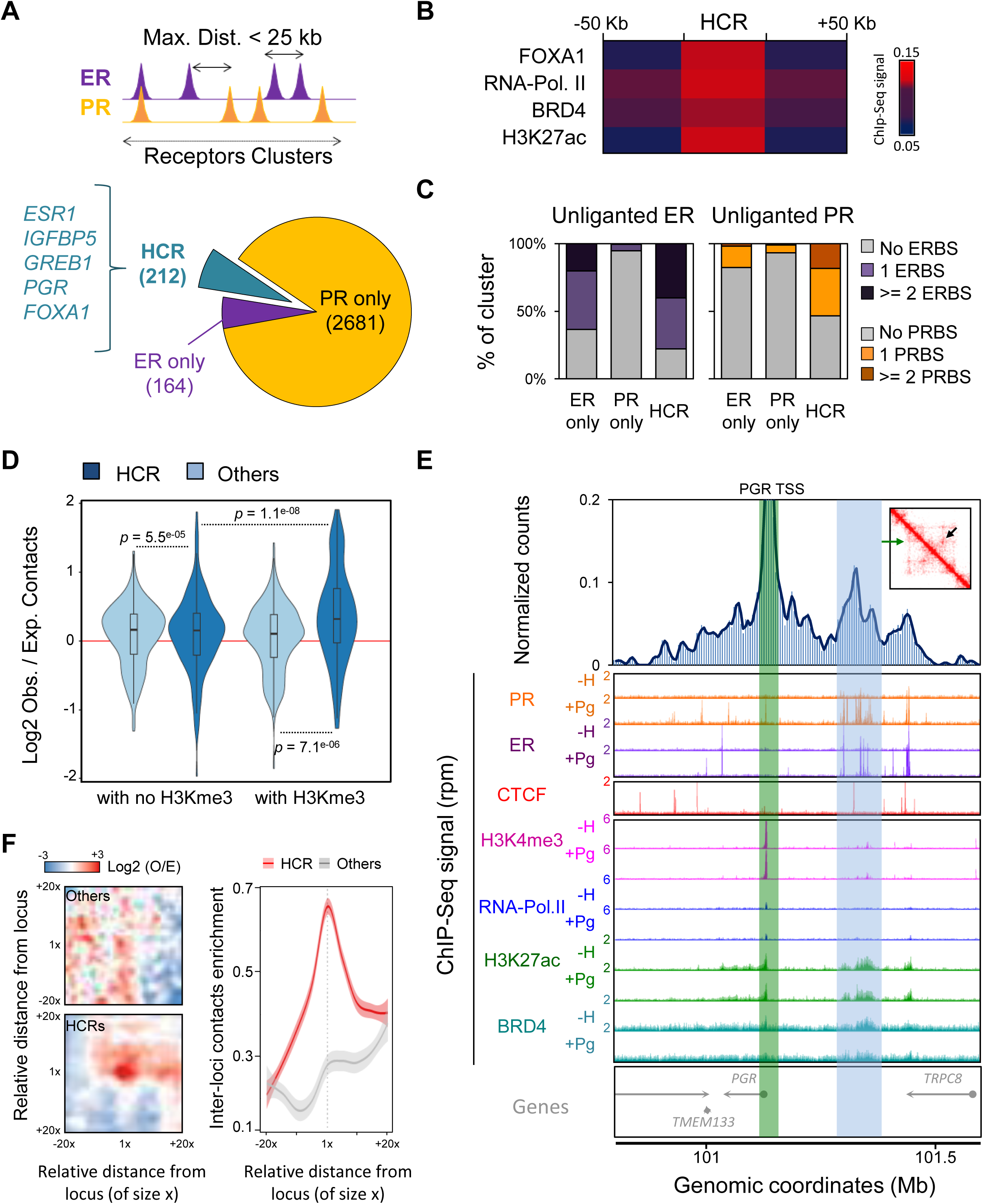
Genome-wide characterization of HCRs. **(A)** Strategy used to identify clusters of receptor binding sites (also see Material and methods section). The pie chart indicates the number of clusters of each type. Examples of genes of interest found in TADs harboring a HCR are shown on the side. **(B)** Average ChIP-Seq signal of specific factors and marks in HCRs and their flanking regions (±50 kb upstream or downstream). **(C)** Percentage of cluster (either ER-predominant, PR-predominant or HCRs) exhibiting 0, 1 or more than 1 binding site(s) for ER (left panel) or PR (right panel) in cells maintained in conditions depleted of steroid hormones (unliganded ER or PR). **(D)** Distribution of observed versus expected intra-TAD interactions established between HCRs or non-HCRs regions (Others) with loci marked or not by H3K4me3. p-values of differences between the different groups are shown (Student t-test). **(E)** *Top panel*: virtual 4C profile at 5 kb resolution using the TSS of *PGR* (highlighted in green) as bait (Chr. 11: 100,800,000-101,600,000). The position of the largest HCR detected in this region is highlighted in blue. The inset shows the corresponding contact matrices obtained in T47D cells grown in absence of hormones. The green arrow shows the position of the TSS of *PGR* and the black arrow points to the loop established between the promoter and the HCR. Panels below correspond to Genome Browser tracks of ChIP-Seq profiles obtained for the different factors and epigenetic marks listed in absence (-H) or presence of Progestin (+Pg) or Estradiol (+E2). Position of protein coding genes and genomic coordinates are shown at the bottom. **(F)** Analysis of the long-range inter-TADs interactions established between HCRs (HCR-HCR) or between random regions of similar sizes and separated by similar distances. On the left panel, X- and Y-axis correspond to the relative genomic coordinates considering the region of interest (of size x) as center. Color scale reflects the observed versus expected contacts (also see Materials and methods section). The graph on the right corresponds to the average (± confidence interval) signal observed at the loci of interest (red: HCR-HCR contacts; grey: RandomRandom contacts).

To determine whether these HCRs corresponded to long-range regulatory elements as observed in the case of the ESR1-TAD, we generated *in situ* Hi-C samples from T47D cells grown in steroid-depleted conditions. We computed the frequencies of contacts of the identified HCRs within their host TADs and observed that they established more frequently intra-TAD interactions with sites marked by H3K4me3 than with other loci (Figure 2D). Besides, HCR-H3Kme3 occured at higher frequency than the contacts between H3K4me3 sites and other loci within the TAD, supporting their role in controlling gene expression (Figure 2D). For example, such interactions were observed between the promoter of *PGR* gene and the HCRs detected in its TAD (Figure 2E). We further used the *in situ* Hi-C data to measure the frequency of contacts between HCRs located > 2 Mb apart within the same chromosome. We observed a significant tendency of long-range HCR-HCR intra-chromosomal contacts as compared to the contacts between random regions of similar sizes separated by similar distances (Figure 2F). These observations point towards a regulatory and structural role of HCRs in the absence of hormones not only at the TAD level but also at higher levels of chromosomal organization.

### Differences in the structural organization of TADs in ER^+^/PR^+^ and ER^−^/PR^−^ breast cells correlate with HCR occupancy by the receptors

Further Analysis of ER and PR ChIP-Seq experiments showed that clustering of ER and PR in regions corresponding to HCRs in T47D also occurred in same regions in MCF-7 despite the very different levels of ER and PR and the limited overlap between binding sites in these breast cancer cell lines (Supplementary Figure 2A). Out of the 212 HCRs identified in T47D cells, 61 (28.7%) were classified as such in MCF-7 cells (PR≥3; ER≥3). For example, the E2-induced binding of ER to the ESR1-TAD HCR at sites already occupied in hormone-depleted conditions was also observed in MCF-7 cells (Supplementary Figure 2B). Upon exposure to Pg, PR also bound to the ESR1-TAD HCR in MCF-7, in this case at sites mainly overlapping with the ER binding sites and different from the PR binding sites observed in T47D (Supplementary Figure 2B). In T47D cells, the binding of ER and PR in the region of clustering occurred frequently at non-overlapping sites (Supplementary Figure 2C). In contrast, there was a higher overlapping between the binding sites of the two receptors in MCF-7 cells within the same regions (Supplementary Figure 2C). Thus, it appears that while ER and PR cluster in qualitatively and quantitatively different ways in T47D and MCF-7 cells, they do so within similar regions.

We then compared the structure of HCR-containing TADs in ER^+^-PR^+^ breast cancer cell lines (T47D, MCF-7, and BT474) and in ER^−^-PR^−^ cells (MCF10A and SKBR3), all grown in absence of hormones (Supplementary Table 1). We observed that regions identified as HCRs in T47D were also engaging long-range interactions between them in the ER^+^-PR^+^ cell lines but not in the ER^−^-PR^−^ ones, or in the ER^−^-PR^−^ lymphoblastoid cell line GM12878 (Figure 3A). These regions also engaged in frequent interactions with sites marked by H3K4me3 within their own TAD in ER^+^-PR^+^ but not in ER^−^-PR^−^ cell lines (Figure 3B). For instance, when we compared the organization of the ESR1-TAD in the different cell lines, we observed structural differences that suggested higher similarities between cells of the same steroid receptor status (Supplementary Figure 3A). The main structural differences corresponded to the HCR interactions, which did not establish contacts with the promoters in ER^−^-PR^−^ cells (Supplementary Figure 3B-C). Similarly, in the absence of hormone, the promoter of *PGR* gene established contacts with HCR in the three ER^+^-PR^+^ cell lines but not in the ER^−^-PR^−^ ones (Figure 3C-E, Supplementary Figure 3D-E). These observations support the notion that HCRs organize the intra-TAD contacts as well as higher levels of genome structure in ER^+^-PR^+^ cells. In addition, they suggest that the binding of the ER and PR in receptor positive cells in the absence of hormone (Figure 2C, Supplementary Figure 2B) might be responsible for the cell specific organization of HCR containing TADs.

**Figure 3:**
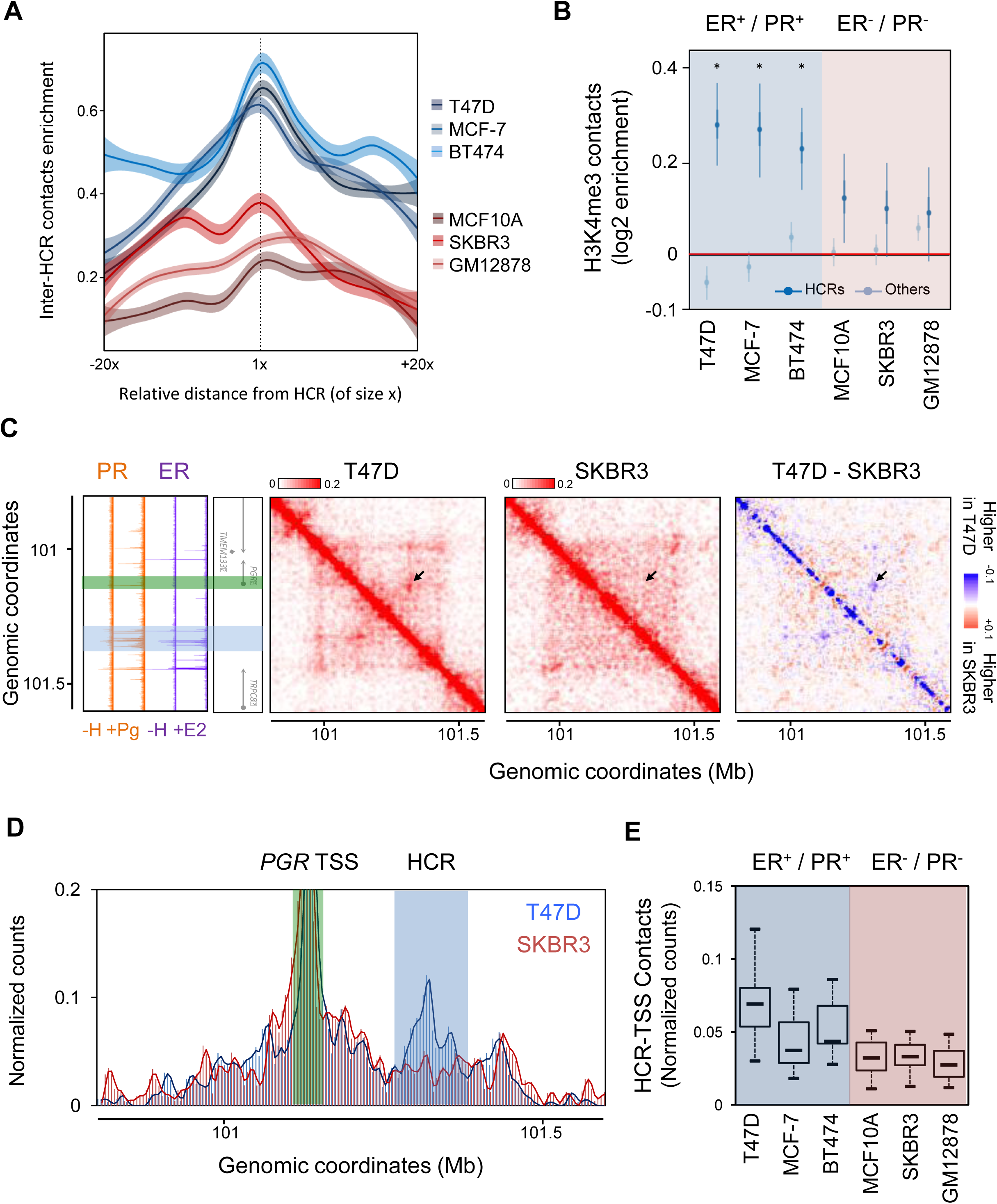
Internal structures of HCR containing TADs correlate with ER/PR status. **(A)** Analysis of long-range inter-TAD interactions established in different cell lines between regions corresponding to the HCRs defined in T47D. The graph shows the average (± confidence interval) of contacts with HCR located at more than 2 Mb using HCRs as in Figure 2F (Red lines: ER^−^-PR^−^ cell lines; Blue lines: ER^+^-PR^+^ cell lines). **(B)** Distributions of observed versus expected intra-TAD interactions established between the HCRs identified in T47D with loci marked by H3K4me3 in the different ER^+^-PR^+^ and ER^−^-PR^−^ cell lines analyzed (dark blue bars). The light blue bars correspond to the observed versus expected intra-TAD interactions established between non-HCR loci with loci marked by H3K4me3 in the same TADs. *p*-values indicated at the top correspond to the pair-wise comparisons of the two distributions for each cell line (paired t-test). **(C)** Normalized *In Situ* Hi-C contact matrices at 5 kb resolution over the PGR-TAD (Chr.11: 100,800,000-101,600,000) obtained in T47D and SKBR3 cells (color scaled according the normalized counts per thousand in the region depicted). The difference between the contact matrices is shown on the right with blue and red corresponding to contacts higher and lower in T47D than in SKBR3, respectively. Panels on the left correspond to Genome Browser tracks (as in Figure 2E) of ChIP-Seq profiles obtained for the different factors listed in the different conditions (−H, +Pg, +E2). Position of protein coding genes and genomic coordinates are shown. Positions of the TSS of *PGR* and its nearby HCR are highlighted in green and blue, respectively. **(D)** Virtual 4C profiles at 5 kb resolution obtained using the promoter of *PGR* as bait in T47D (blue) or SKBR3 (red). The positions of the TSS of *PGR* and its nearby HCR are highlighted in green and blue, respectively. **(E)** Distribution of contacts between genomic bins within the HCR with the TSS of *PGR.*

### Hormone independent binding of ER at HCRs participates in the maintenance of a functional TAD organization

The binding sites for CTCF within the TADs described above was highly similar in ER^+^-PR^+^ and ER^−^-PR^−^ cells (Supplementary Figure 3F). Moreover, the functional interactions of HCRs were observed in cells expressing ER and PR in absence of steroids. We therefore wondered whether the fraction of bound unliganded receptors (Figure 2C) could directly participate in the maintenance of such organization. To test this hypothesis, we knocked-down ER expression in T47D cells by transient transfection with siRNA against ERα (siRNA-ER). In two independent experiments, a mild decrease (45%) of the levels of ER in siRNA-ER transfected cells was accompanied by a concomitant decrease of the level of PR transcripts in hormone depleted conditions (Figure 4A). This was accompanied by a decrease of the interactions between the *PGR* promoter and HCRs in siRNA-ER transfected cells as compared to cells transfected with scramble siRNA (siRNA-Control; Figure 4B). Similarly, we observed a decrease in the interactions between HCRs and promoters within the ESR1-TAD (Figure 4C). Comparable trends were observed comparing T47D cells where PR levels were knocked-down by shRNA, which leads to a concomitant decrease of ER levels (Verde et al. 2017*). *In situ* Hi-C performed in shEmpty or shPR cells also suggested a differential organization of the ESR1- and the PGR-TADs, with the interactions between the HCRs and the promoters partially disrupted in shPR cells (Supplementary Figure 4). Although limited due to the modest decrease of receptor levels reached in the knock-down experiments, these observations support the hypothesis that steroid receptors bound to chromatin in absence of ligand serve to organize and maintain HCR interactions prior to hormone exposure.

**Figure 4:**
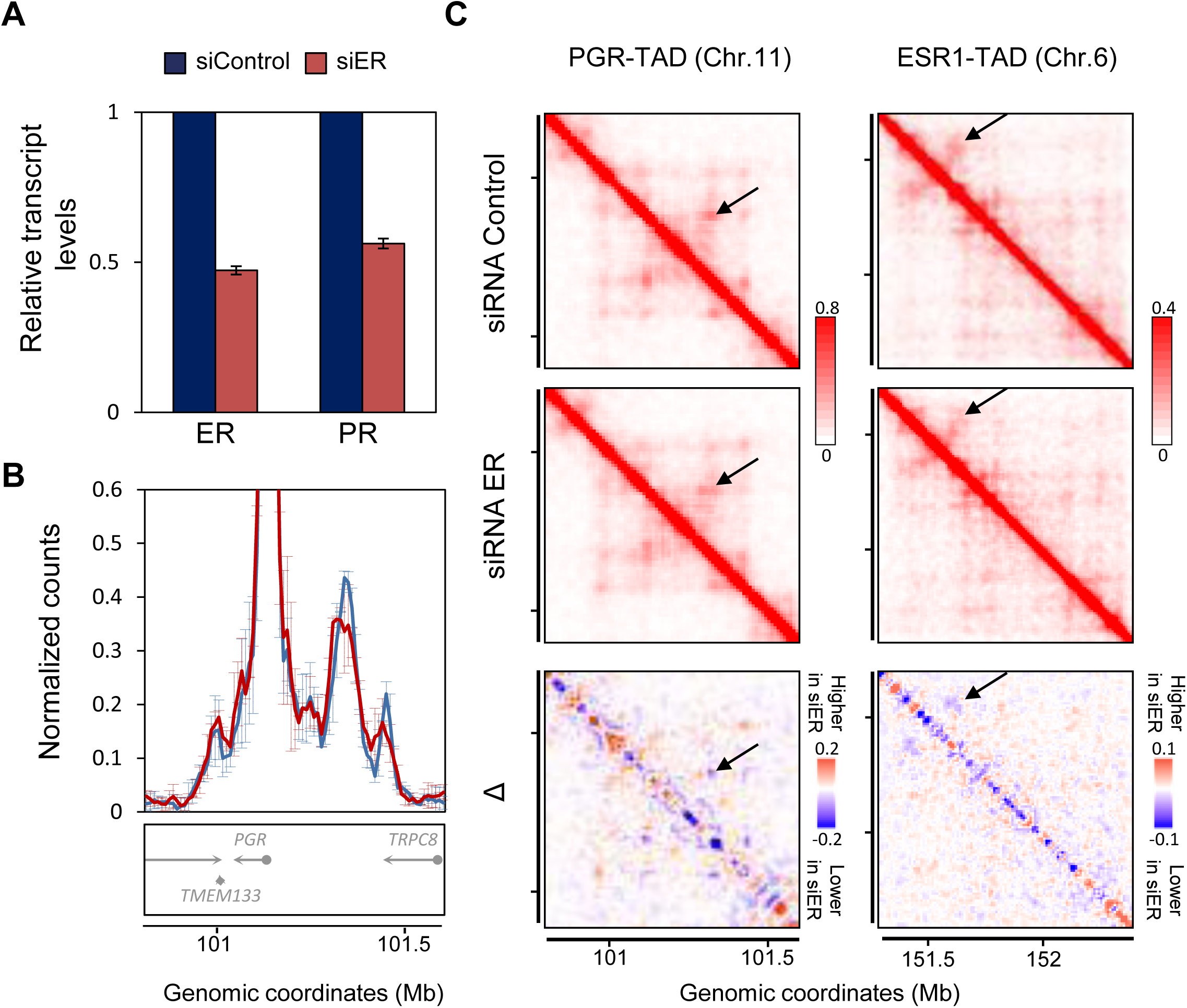
**(A)** Relative levels of ER and PR transcripts in T47D cells transfected with scramble siRNA (siRNA-Control) or siRNA against ER (siRNA-ER). **(B)** Virtual 4C profiles at 10 kb resolution using the TSS of *PGR* as bait in T47D siRNA-Control cells (blue line) or in siRNA-ER cells (red line) grown in the absence of hormones (average ± SD of two independent transient transfections). **(C)** Normalised *In Situ* Hi-C contact matrices at 10 kb resolution over the ESR1-TAD (Chr.6: 151,300,000-152,400,000; left panels) and PGR-TAD (Chr.11: 100,800,000-101,600,000; right panels) obtained in T47D siRNA-Control cells (top) or in siRNA-ER cells (middle) grown in the absence of hormones. The difference between the contact matrices is shown at the bottom with blue and red corresponding to contacts higher and lower in T47D siRNA-Control than in T47D siRNA-ER, respectively.

### HCR-containing TADs are dynamically restructured upon exposure to hormones

To further interrogate the functional role of the identified HCRs, we generated *in situ* Hi-C from T47D cells exposed to Pg or E2 for 30 or 180 minutes (Supplementary Table 1). The long-range inter-TAD interactions between HCRs were mainly maintained upon exposure to the two hormones (Figure 5A). When analyzing the Pg-induced modifications of enhancer-associated marks at HCRs in T47D cells, we further noticed that two distinct populations of HCRs could be identified based on hormone-induced changes in BRD4 signal: 131 loci showed enhanced recruitment of BRD4 upon exposure to Pg, correlating with increased H3K27ac and RNA Pol. II levels. We refer to them as HCR^+^ (Figure 5B). In contrast, exposure to Pg decreased the level of these marks in the remaining 81 HCRs that we refer to as HCR^−^ (Figure 5B). We thus calculated the global transcriptional changes that occur within HCR^+^ or HCR^−^ containing TADs in response to exposure to Pg or E2, and observed that HCR^-^ were associated with E2-stimulated but Pg-repressed activities, as in the case of the ESR1- or PGR-TADs. In contrast, both Pg and E2 appeared to enhance the transcriptional activity of TADs containing HCR^+^ (Figure 5C). These opposite effects of E2 and Pg on HCR^−^ were confirmed when considering more specifically the enrichment of significantly hormone-regulated protein-coding genes located within HCR-containing TADs (Figure 5D).

**Figure 5:**
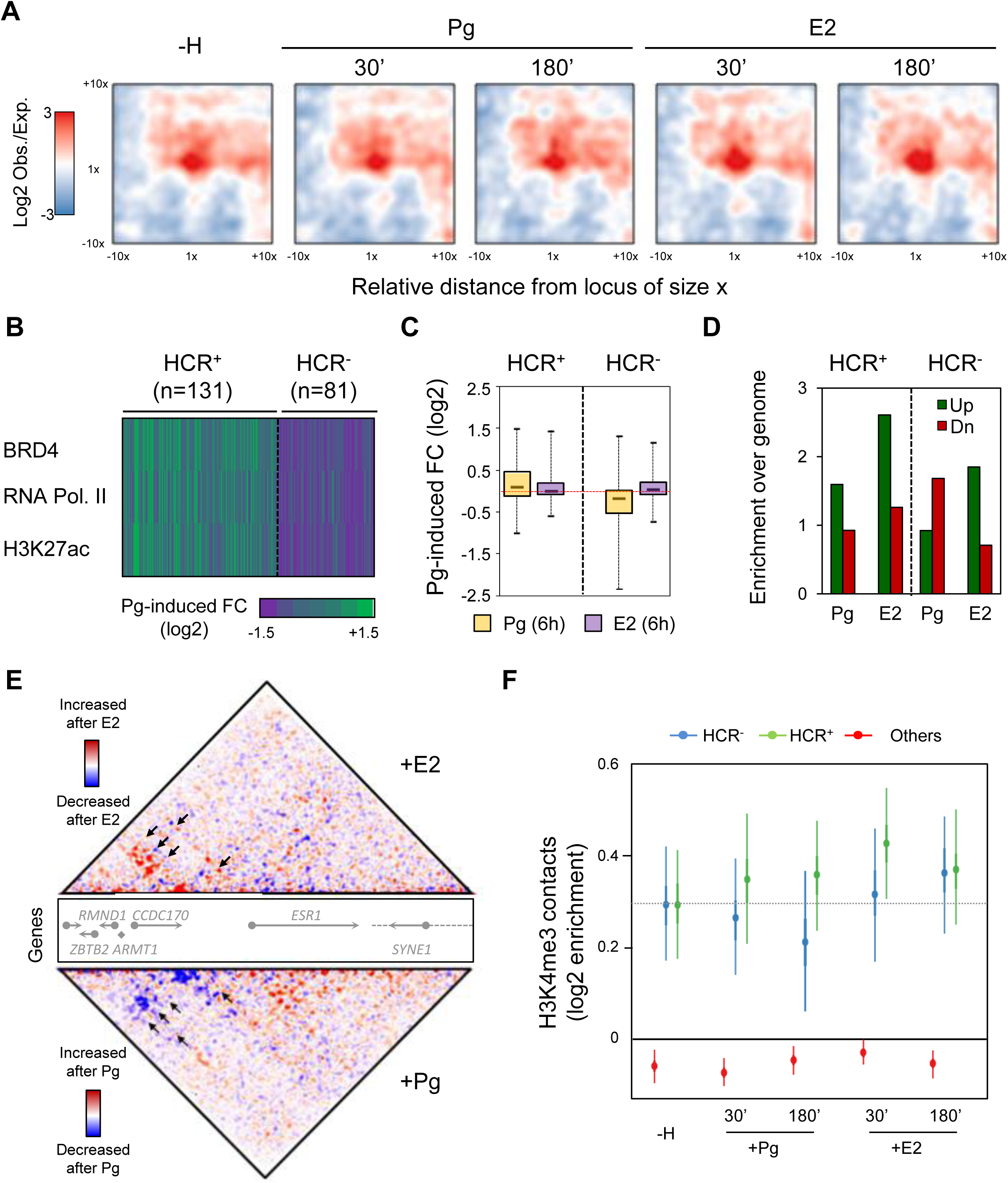
Hormones dynamically modify HCR-containing TAD structure and interactions. **(A)** Analysis of the long-range inter-TADs interactions established between HCRs before (as in Figure 2F) or after 30 and 180 minutes exposure to Pg or E2. **(B)** Fold change in the levels of BRD4, RNA Pol. II and H3K27ac after 30 minutes of treatment with Pg (green: increased signal after hormone, purple: lower signal after exposure to the hormone). Depending on the increase or decrease in BRD4 signal after exposure to Pg, we classified HCRs as HCR^−^ or HCR^+^, respectively. **(C)** For each TAD containing a HCR^−^ or a HCR^+^, global changes of expression after 6h of treatment with Pg or E2 were computed. Boxplots show the distributions of these Pg- or E2-induced fold-changes per TADs. **(D)** The number of significantly Up- (green) or Down- (red) regulated genes by progesterone or E2 in TADs containing HCR^+^ or HCR^−^ were compared to the expected proportion of the genome. Histograms show the levels of enrichment of E2- or Pg-Up- (green) or E2- or Pg-Down-regulated (red) genes within TADs containing HCR^+^ or HCR^−^. **(E)** Differential analysis of the contacts obtained from Capture-C experiment on the ESR1-TAD before and after 60 minutes exposure to E2 (top panel) or Pg (bottom panel). Contacts more frequent after hormone exposure are highlighted in red whereas contacts decreasing after hormone are labeled in blue on the heatmaps. **(F)** Enrichment of intra-TAD contacts between either HCR^+^ (green) or HCR^-^ (blue) and H3K4me3 peaks before or after exposure to 30 and 180 minutes of E2 or Pg.

To explore the associated structural changes, we analyzed at high-resolution how the organization of the HCR^-^ containing ESR1-TAD was modified upon hormone treatment using the enrichment probes described in Figure 1A in T47D cells exposed or not to Pg or E2 for 60 minutes (Supplementary Table 1). In independent experiments of Capture-3C and Capture-Hi-C, we observed opposite effects of the two hormones on the internal organization of the ESR1 - TAD (Supplementary Figure 5A). We pooled these capture datasets and performed differential analysis of the contact maps before/after exposure to the hormones. These comparisons demonstrated differential hormone-induced reorganization of the HCR contacts: ESR1-TAD HCR^−^ interactions with promoters increased upon E2 exposure but decreased in response to Pg (Figure 5E, top panel and bottom panels, respectively). These opposite effects were confirmed after 180 minutes of exposure to the hormones (Supplementary Figure 5A). Thus, the ESR1-TAD HCR^−^ acts as an enhancer in absence of hormone when ER is already bound in a ligand-independent manner and its activity is further enhanced upon exposure to E2. In contrast, upon exposure to Pg binding of PR to the HCR^−^ decreased its activity and destabilized the TAD structure. Genome-wide analysis of *in situ* Hi-C performed in T47D cells exposed to Pg or E2 for 30 or 180 minutes confirmed that the two hormones enhance the interactions of the regulatory regions with promoters in the case of HCR^+^ containing TADs (Figure 5F). In contrast, whereas E2 also enhances the HCR-promoters interactions within HCR^−^ containing TADs, Pg contributes to the destabilization of the interactions engaged by HCR^−^ (Figure 5F). For example, as observed in the case of the ESR1-TAD, the interactions engaged by the HCR^−^ located within the PGR-TAD are enhanced upon exposure to E2 but decreased in response to Pg in correlation with the opposite activities of the hormones on the expression of the PGR gene (Supplementary Figure 5B). Together these results suggest that steroid receptors bound in absence of their cognate ligand can maintain specific interactions of large regulatory regions and that these interactions are dynamically and, in some cases, differentially modify upon further binding of the ER and PR receptors in response to hormones.

## Discussion

Here, we report the analysis at high resolution of the internal structure of the ESR1-TAD, which is coordinately regulated by steroid hormones in T47D cells. We found that genes within this TAD are organized around a 90 kb region of clustered binding sites for ER and PR. In the absence of hormones, this Hormone Control Region (HCR) already contacts the promoters of all genes within the TAD maintaining a basal expression of the resident genes. Genome-wide we identified more than 200 of such putative HCRs, which, as previously described for other locus control regions (Fraser and Grosveld 1998; Li et al. 2002), organize long-range interactions with promoters within their respective TAD, as well as inter-TAD interactions between them. HCRs are characterized by enhancer-related marks and recruitment of transcription associated proteins (RNA Pol. II and BRD4) and are frequently linked to genes involved in the identity of breast epithelial cells as in the case of the *ESR1*, *PGR* and *FOXA1* genes. They could therefore correspond to previously described stretch-enhancers (Parker et al. 2013) or super-enhancers (Whyte et al. 2013), as in the case of the ESR1-TAD HCR (Hnisz et al. 2015). Indeed, it has been previously proposed that super-enhancers could organize modules of integration of different signaling pathways and that perturbations of the activity of these platforms can lead to major changes in the expression of the dependent genes (Hnisz et al. 2015). In addition to support these observations, our results further demonstrate that the activity of the HCRs is highly dynamic and modular. Only 27 out of the 61 common MCF-7 and T47D HCRs have been previously classified as super-enhancers in MCF-7 (Hnisz et al. 2015). This limited overlap, potentially due to lower levels of acetylated H3K27 at HCRs than at the previously identified super-enhancers in MCF-7 (Hnisz et al. 2015), suggests that HCRs correspond to a specialized class of modular regulatory elements. In particular, we identified 81 of these regions where binding of the ER and PR upon exposure to the hormones in T47D cells leads to opposite effects on gene expression as well as in their contacts with the rest of the TAD. We therefore suggest that these regions could act as enhancers or silencers depending on the signal received. This signal-dependent modulation of HRC activity argues that regulatory regions not only integrate various synergistic signals but that different modules can cross-talk between them in a more complex manner. Indeed, it appears that HCR are complex regulatory elements spanning several tens of kb within which the two receptors bind at distinct sites resulting in a fine-tuned activity of the target genes. Depending on the HCR considered, Pg and E2 can have similar effects, as exemplified by the increased levels of enhancer associated marks and activity in response of the HCR^+^ to E2 or Pg. In contrast, in the case of the HCR^−^, the two hormones have opposite effects on gene expression and chromatin organization. These observations therefore suggest more complex relations between estrogens and progestins signaling pathways than previously described. Indeed, PR has been proposed to be an important mediator or ER activity by modifying the landscape of binding of ER (Mohammed et al. 2015). Our observations further show that ER and PR can bind within large genomic regulatory regions, either on overlapping or nonoverlapping sites, where the receptors can have distinct activities on gene expression.

Cell specific intra-TAD organization, reflected by the establishment of cell specific sub-domains or chromosome neighborhoods (Phillips-Cremins et al. 2013; Dowen et al. 2014), has been proposed to depend on differential binding of architectural proteins like CTCF and cohesins together with subunits of the Mediator complex (Phillips-Cremins et al. 2013). The binding of CTCF remains largely similar, even between cells of different origins and of divergent ER/PR status. Differential binding of this architectural protein might therefore not be sufficient to explain the observed structural differences. In contrast, cells with divergent ER/PR status exhibit different intra-TAD organization, which correlate with the expression of the receptors and their binding at HCRs in absence of hormones. Lowering the levels of receptors in ER^+^-PR^+^ cells leads to a destabilization of the TAD structure, pointing towards direct roles of the nuclear receptors in the organization of intra-TAD chromatin folding in the absence of hormones. It has been previously proposed that transcription factors act on a structurally pre-organized context with loops between promoters and enhancers already primed prior to activation of the transcription factors (Hakim et al. 2009; Jin et al. 2013). Although the clustering of receptors is more evident after exposure to their cognate steroid hormones, our results suggest that a large fraction of HCRs is already partially occupied by the receptors, in particular by ER, in hormone-depleted conditions. This fraction of sites occupied by unliganded receptors might be therefore sufficient and necessary to maintain the apparent pre-established chromatin folding structure, facilitating further response. The fact that HCRs not only organize the 3D structure within TADs but also appear to engage long-range interactions between them suggest the existence of specific nuclear hubs where specific TADs could be enriched. Although probably highly stochastic within the cell population, these hubs of regulatory regions might serve as a reservoir of factors to facilitate transcription in a permissive environment. They may correspond specialized transcription factories (Xu and Cook 2008) or to active hubs previously described for other steroid receptors (Hakim et al. 2011; Kuznetsova et al. 2015). Other transcription factors, as for example STAT transcription factors, have also been proposed to play a direct functional role in organizing the genome of T cells by leading to the spatial clustering of regulatory regions in specific hubs (Hakim et al. 2013).

Interactions between HCRs and promoters are further modified upon exposure to the hormones and increased binding of the receptors, supporting a direct role of the activated transcription factors in modifying these contacts, either by reinforcing or destabilizing them. Interestingly, these regulatory regions can establish contacts with sites that lack actual binding sites of the receptors (for instance ERS1-TAD HCR with promoters Figure 1C), suggesting that the regulatory loops might be established between receptors or receptor associated factors and other transcription factors and/or structural proteins of chromatin. Further studies will be required to properly understand how nuclear receptors facilitate the maintenance of these long-range intra-TAD contacts. In particular, it remains to be investigated whether additional transcription factors, as for example FOXA1 (Figure 2B), also participate in establishing and/or maintaining the 3D structure of HCR-containing TADs. In addition, Estrogens and progestins rapidly activate protein kinases (Vicent et al. 2006) and PARP1 (Wright et al. 2012) leading to the synthesis of nuclear ATP, which is required for the response to estrogen and progestins (Wright et al. 2016). These pathways could have important roles in the structuration of the genome, which also remains to be investigated.

In summary, we propose that, in absence of steroid hormones, a fraction of unligaded receptors participate in the maintenance of a TAD structure that is permissive for the regulation of gene expression. Upon exposure to the hormones, further binding of the receptors can enhance or, in contrast, destabilize the interactions established between HCR and promoters leading to adequate response to the signals. Together, our results support a relevant role of cell specific transcription factors in shaping the genome in a way that contributes to the integration of the cell response to external cues. It will now be important to determine how these activities are accomplished; in particular, which are the epigenetic factors that determine the positive and negative effects of receptors recruitment to the DNA.

## Material and methods

### Cell lines and culture conditions

T47D, MCF-7, BT474, MCFOA, SKBR3 were routinely grown in their optimal medium according the ATCC recommendations. Stable T47D clones expressing shRNA empty or shRNA against PR are described elsewhere (Verde et al., 2017*). For RNA interference experiments, T47D cells were transiently transfected as previously described (Nacht et al. 2016) with scramble siRNA (siRNA-Control) or siRNA directed against ERα (siRNA-ER; sc-29305, Santa-Cruz Biotechnology). For the experiments cells were grown 48 hours in medium without phenol red complemented with 10% charchoal-dextran treated Fetal Bovine Serum (which corresponds to steroid depleted condition). After further synchronization in G0/G1 by overnight culture in medium without phenol red in absence of FBS, cells were harvested (steroid depleted (−H) condition). In the case of hormonal treatments, cells were further treated for the times QQ indicated with 10^−8^ M Estradiol (E2) or with 10^−8^ M of the progestin analogue R5020 (Pg).

### Chromatin immunoprecipitation (ChIP) and sequencing analysis

ChIP-seq experiments in T47D cells for H3K4me3, RNA-polymerase II, PR, FOXA1 and CTCF were described previously (Ballare et al. 2013; Le Dily et al. 2014; Nacht et al. 2016). Additional ChIP-seq experiments for BRD4 (, ERα (HC20, Santa-Cruz Biotechnology) and H3K27ac (ab4729, Abcam) were performed as described previously (Nacht et al. 2016). After library preparation and sequencing on a HiSeq2000, reads were trimmed and processed by aligning to the reference human genome (GRCh38/hg38) using BWA (Li and Durbin 2009). Only uniquely aligned reads were conserved. ChIP-seq signals were normalized for sequencing depth (expressed as reads per milions – rpm). and we identified binding sites (or peaks) using MACS2 (Zhang et al. 2008).

### Genome-wide identification and classification of receptors clusters

Genomic clusters of ER and/or PR were determined by using ChIP-Seq data of ER and PR in T47D exposed 60 minutes to E2 or Pg, respectively. Binding sites for any of the two receptors (q-value < 10^−6^) separated by less than 25 Kb were aggregated. Receptors clusters were classified as follow: ER-clusters were regions with ER binding sites ≥ 3 and PR binding sites < 3; PR-clusters were regions with PR binding sites ≥ 3 and ER binding sites < 3; HCR correspond to regions with ER binding sites ≥ 3 and PR binding sites ≥ 3.

### In situ Hi-C library preparation

*In situ* Hi-C experiments were performed as previously described (Rao et al. 2014) with some modifications. Adherent cells were directly cross-linked on the plates with 1% formaldehyde for 10 minutes at room temperature. After addition of glycine (125 mM final) to stop the reaction, cells were washed with PBS and recovered by scrapping. Cross-linked cells were incubated 30 minutes on ice in 3C lysis Buffer (10 mM Tris-HCl pH=8, 10 mM NaCl, 0.2% NP40, 1X anti-protease cocktail), centrifuged 5 minutes at 3,000 rpm and resuspended in 190 uL of NEBuffer2 1X (New England Biolabs – NEB). 10μL mL of 10% SDS were added and cells were incubated for 10 minutes at 65°C. After addition of Triton X-100 and 15 minutes incubation at 37°C, nuclei were centrifuged 5 minutes at 3,000 rpm and resuspended in 300 uL of NEBuffer2 1X. Digestion was performed overnight using 400 U MboI restriction enzyme (NEB). To fill-in the generated ends with biotinylated-dATP, nuclei were pelleted and resuspended in fresh 1x repair buffer 1.5 μL of 10 mM dCTP;1.5 μL of 10 mM dGTP; 1.5 μL of 10 mM dTTP; 37.5 μL of 0.4 mM Biotin-dATP; 50U of DNA Polymerase I Large (Klenow) fragment in 300 uL NEBuffer2 1X). After 45 minutes incubation at 37°C, nuclei were centrifuged 5 minutes at 3,000 rpm and ligation was performed 4 hours at 16°C using 10,000 cohesive end units of T4 DNA ligase (NEB) in 1.2 mL of ligation buffer 120 μL of 10X T4 DNA Ligase Buffer; 100 μL of 10% Triton X-100; 12 μL of 10 mg/mL BSA; 963 μL of H2O). After reversion of the cross-link, DNA was purified by phenol extraction and EtOH precipitation. Purified DNA was sonicated to obtain fragments of an average size of 300-400 bp using a Bioruptor Pico (Diagenode; 8 cycles; 20 s on and 60 s off). 3 μg of sonicated DNA was used for library preparation. Briefly, biotinylated DNA was pulled down using 20 uL of Dynabeads Myone T1 streptavidine beads in Binding Buffer (5 mM Tris-HCl pH7.5; 0.5 mM EDTA; 1M NaCl). End-repair and A-tailing were performed on beads using NEBnext library preparation end-repair and A-tailing modules (NEB). Illumina adaptors were ligated and libraries were amplified by 8 cycles of PCR. Resulting Hi-C libraries were first controlled for quality by low sequencing depth on a NextSeq500 prior to higher sequencing depth on HiSeq2000. Supplementary Table 1 summarizes the number of reads sequenced for each of the datasets presented in this study.

### Capture-3C and Capture-Hi-C

3C were performed according the protocol described above for the Hi-C omitting the biotin-labeling step. 120 mer biotinylated RNA probes covering the region Chr.6: 149’450’000-153’750’000 (purchased from Agilent) were designed using the SureDesign software with the following parameters: density: 2X, masking: moderately stringent and boosting: balanced. 3C samples were sonicated to an average size of 300-400 bp and pre-capture libraries were generated using the SureSelect kit according the manufacturer’s instructions. Hybridization and post-capture library amplification were performed according the manufacturer instructions (Agilent). In the case of Capture-Hi-C experiments, Hi-C libraries, generated according the *in situ* protocol described above, were used as pre-capture libraries for the hybridization step.

### Normalization of Hi-C matrices, differential analysis and virtual 4C

Hi-C data were processed using an in-house pipeline based on TADbit (Serra et al. 2017). Reads were mapped according to a fragment-based strategy: each side of the sequenced read was mapped in full length to the reference genome Human Dec. 2013 (GRCh38/hg38). In the case reads were not mapped, they were split when intra-read ligation sites were found and individual split read fragments were then mapped independently. We used the TADbit filtering module to remove non-informative contacts and to create contact matrices as previously described (Serra et al. 2017). PCR duplicates were removed and the Hi-C filters applied corresponded to potential non-digested fragments (extradandling ends), non-ligated fragments (dandling-ends), self-circles and random breaks (Supplementary Table 1 summarizes the number of reads mapped and the number of valid pairs used to generate the matrices for the different samples used in this study). The matrices obtained were normalized for normalized for sequencing depth and genomic biases using OneD (Vidal et al. 2017*) and were further smoothed using a focal (moving window) average of one bin. *In situ* Hi-C data for GM12878 cells were obtained from Rao et al. (2014) and reprocessed as mentioned above. In Figure 1A and 1B, normalized reads are expressed as normalized counts per million of genome-wide valid pairs. In the rest of the analysis, since the cell lines used are aneuploidy and present different number of copies of the chromosomes analyzed, the matrices obtained in the different cell lines or conditions were further normalized for local coverage within the region (expressed as normalized counts per thousands within the region) without any correction for the diagonal decay. For differential analysis, the resulting normalized matrices were directly subtracted from each other. To note, MCF-7 cells have a micro-amplification of a genomic segment of approximately 100 kb, which overlaps the HCR of ESR1-TAD, impeding proper comparison with the other cell lines in this particular region (Supplementary Figure 3A-C).Virtual 4C plots were generated from these normalized matrices and correspond to histogram representation of the lines of the matrices containing the baits (therefore expressed as counts per thousand of normalized reads within the region depicted).

### RNA-Seq

Replicate of RNA-Seq experiments were performed in T47D cells treated or not with 10^−8^ M R5020 or 10^−8^ M E2 during 6 hours as described previously (Le Dily et al. 2014). Together with previously published RNA-Seq replicates in these cells (Le Dily et al. 2014), paired end reads were (re)mapped to the hg38 assembly version of the human genome with GENCODE’s V24 annotation.

### Hormone-induced transcriptional changes and enrichment of regulated genes in HCR-containing TADs

A consensus list of TAD coordinates was generated based on the *in situ* Hi-C datasets from T47D cells with the assumption that the majority of the boundaries are comparable between cell types and upon different conditions of culture (Dixon et al. 2012; Le Dily et al. 2014). The HCRs identified in T47D were assigned to a single TAD and TADs were classified as HCR^+^ or HCR^−^ containing. The total RNA-Seq rpm per TAD in cells treated or not with hormones was computed and we calculated the ratio of changes per TAD as previously described (Le Dily et al. 2014). In addition, we computed the total number of genes significantly regulated by the hormones (p-value < 0.05; −15 < FC > 15) within HCR-containing TADs and compare this observed number to the one expected based on the genome-wide proportions of responsive genes.

### Intra-TAD contacts between HCR and H3K4me3 sites

Using Hi-C matrices at 5 kb resolution, we focused on TADs containing HCRs. Each bin was labeled as part of a HCR or marked by H3K4me3 peaks or others if they did not belong to previous types. Then we gathered the observed contacts between the different types of bins within their TAD and computed expected contacts frequencies based on the genomic distance that separate each pair (the expected distance decay was calculated excluding entries outside TADs). Results are expressed as log2 of the ratio observed on expected frequencies of contacts.

### Inter-TAD contacts between HCR

For each intra-chromosomal pairs of HCR separated by more than 2 Mb, local contact matrices centered on both HCRs were generated. Since HCRs have different sizes, the matrices were generated considering bins of the size × of the HCR and extended upstream and downstream by 20 bins of size × (relative distance to HCR). The observed contacts obtained were corrected for the expected contacts in the case of random regions of similar sizes and separated by the same genomic distance. For each cell line, we excluded the local matrices of regions presenting internal copy number variations in the regions considered. The matrices were smoothed using a focal (moving window) average of one bin and cumulated to generate the meta-contact matrices.

## Acknowledgments

We thank the CRG Ultra-sequencing Facility for technical support and all members of the Chromatin and Gene Expression group for helpful discussions. We acknowledge the members of the 4DGenome project (CRG and CNAG-CRG, Barcelona), notably Thomas Graf, Marc A. Marti-Renom and Guillaume Filion for their helpful comments on the manuscript. We thank Dr. Cheng-Ming Chiang (UT Southwestern Medical center) for kindly providing antibody against BRD4. We received funding from the European Research Council under the European Union’s Seventh Framework Programme (FP7/2007-2013)/ERC Synergy grant agreement 609989 (4DGenome). The content of this manuscript reflects only the author’s views and the Union is not liable for any use that may be made of the information contained therein. We acknowledge support of the Spanish Ministry of Economy and Competitiveness, ‘Centro de Excelencia Severo Ochoa 2013-2017’ and Plan Nacional (SAF2016-75006-P), as well as support of the CERCA Programme / Generalitat de Catalunya.

## Author Contributions

F.L.D. and M.B. designed the study; F.L.D. performed most of the experiments with the help of Y.C.; G.P.V., S.N. and P.S. performed ChIP-Seq experiments; F.L.D. and E.V. carried out the data analysis with contributions of J.Q.; G.V. isolated the shPR cells; All authors discussed the results; F.L.D. and M.B. wrote the manuscript.

## Le Dily et al. - Supplementary Information

**Supplementary Table 1:** Summary of 3C-derived datasets

| Experiment | Sample | Sequenced Read pairs | Valid Read pairs | Raw counts in depicted regions |  |
| --- | --- | --- | --- | --- | --- |
|  |  |  |  | ESR1-TAD | PGR-TAD |
| In Situ Hi-C | BT474 | 196,802,667 | 126,400,945 | 23,014 | 51,322 |
|  | MCF10A | 306,018,506 | 187,513,575 | 48,500 | 31,102 |
|  | MCF7 | 320,056,659 | 166,607,495 | 43,758 | 12,056 |
|  | SKBR3 | 205,997,414 | 156,557,711 | 39,534 | 24,338 |
|  | GM (Rao et al., 2014) | 190,079,469 | 169,692,167 | 45,404 | 20,586 |
|  | T47D T0 | 189,967,400 | 157,390,317 | 36,962 | 32,592 |
|  | T47D R30 | 179,252,416 | 145,811,017 | 34,576 | 30,150 |
|  | T47D R180 | 192,668,851 | 158,516,232 | 40,888 | 34,610 |
|  | T47D E30 | 249,799,510 | 193,968,773 | 51,230 | 47,276 |
|  | T47D E180 | 197,859,885 | 161,304,275 | 39,310 | 34,986 |
|  | T47D shEmpty | 204,764,799 | 128,911,715 | 28,984 | 23,990 |
|  | T47D shPR | 166,975,353 | 114,011,263 | 23,224 | 18,712 |
|  | T47D siRNA Control rep.1 | 206,292,967 | 116,641,257 | 26,752 | 26,378 |
|  | T47D siRNA Control rep.2 | 205,462,437 | 109,936,278 | 28,298 | 19,762 |
|  | T47D siRNA ER rep.1 | 209,173,554 | 118,085,021 | 33,092 | 24,388 |
|  | T47D siRNA ER rep.2 | 204,518,224 | 111,152,397 | 34,296 | 19,334 |
| Capture-3C | T47D T0 | 325,166,775 | 57,414,832 | 1,564,728 | ND |
|  | T47D R60 | 283,013,228 | 43,446,033 | 1,594,758 | ND |
|  | T47D R180 | 281,176,954 | 47,767,277 | 1,953,130 | ND |
|  | T47D E60 | 282,812,094 | 26,517,206 | 935,620 | ND |
| Capture-HIC | T47D T0 | 45,857,588 | 18,997,544 | 246,450 | ND |
|  | T47D R60 | 32,313,052 | 13,824,138 | 217,086 | ND |
|  | T47D R180 | 22,555,505 | 8,952,798 | 101,310 | ND |
|  | T47D E60 | 23,576,054 | 9,922,720 | 168,352 | ND |
|  | T47D E180 | 13,505,153 | 5,653,815 | 111,758 | ND |

\*Valid pairs after filtering of: PCR duplicates, self-circle, dandling-ends, random breaks and errors.

**Supplementary figure 1 (related to Figure 1):**
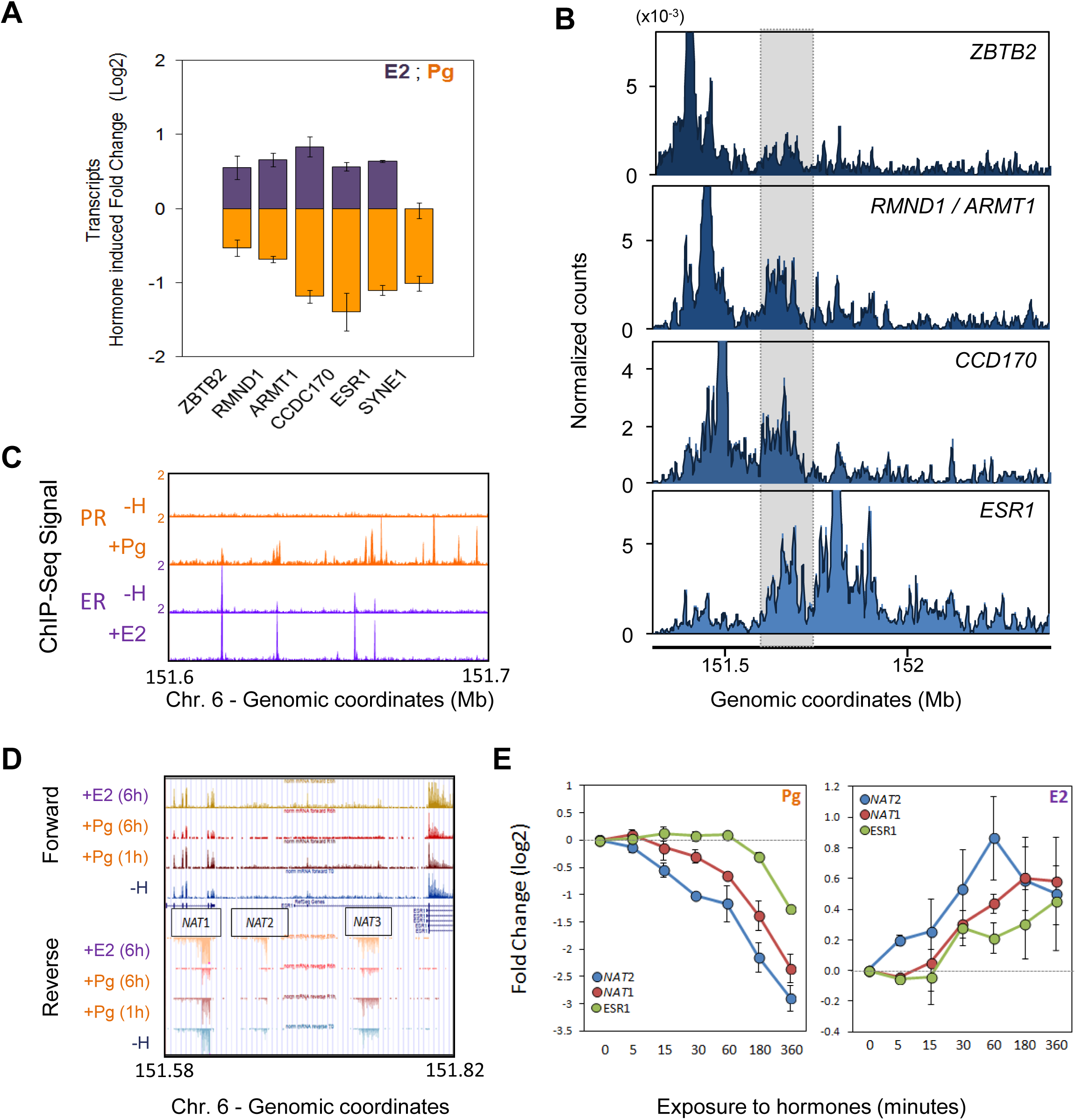
**(A)** Log2 fold change of expression of protein-coding genes located in the ESR1-TAD after 6 hours of exposure to Pg (Orange) or E2 (Purple). The plot shows the average ± SEM expression on 3 independent RNA-Seq experiments. **(B)** Virtual 4C profiles at 2 kb resolution using the TSS of genes within the ESR1-TAD as baits. Data were obtained from T47D cells grown in absence of hormones. **(C)** Magnification of the Genome Browser tracks from Figure 1C: ChIP-Seq profiles obtained for PR and ER in absence (−H) or presence of Pg (+Pg) or E2 (+E2) within the region Chr.6: 151,600,000-151,700,000. **(D)** RNA-Seq profiles obtained in T47D grown in absence of hormones (−H) or exposed 60 or 360 minutes to Pg (+Pg) or 360 minutes to E2 (+E2) showing the expression of 3 distinct non-annotated transcripts (*NAT1*, *NAT2* and *NAT3*) within the region. **(E)** Quantitative RT-PCR showing the kinetics of changes of *NAT1*, *NAT2* and *ESR1* after exposure to Pg (left panel) or E2 (right panel) showing that the non-annotated transcripts comprised in the ESR1-TAD HCR are coordinately regulated by the hormones similarly to the protein coding genes. Plots correspond to the average (± SEM) from two independent kinetics.

**Supplementary figure 2 (related to Figure 3):**
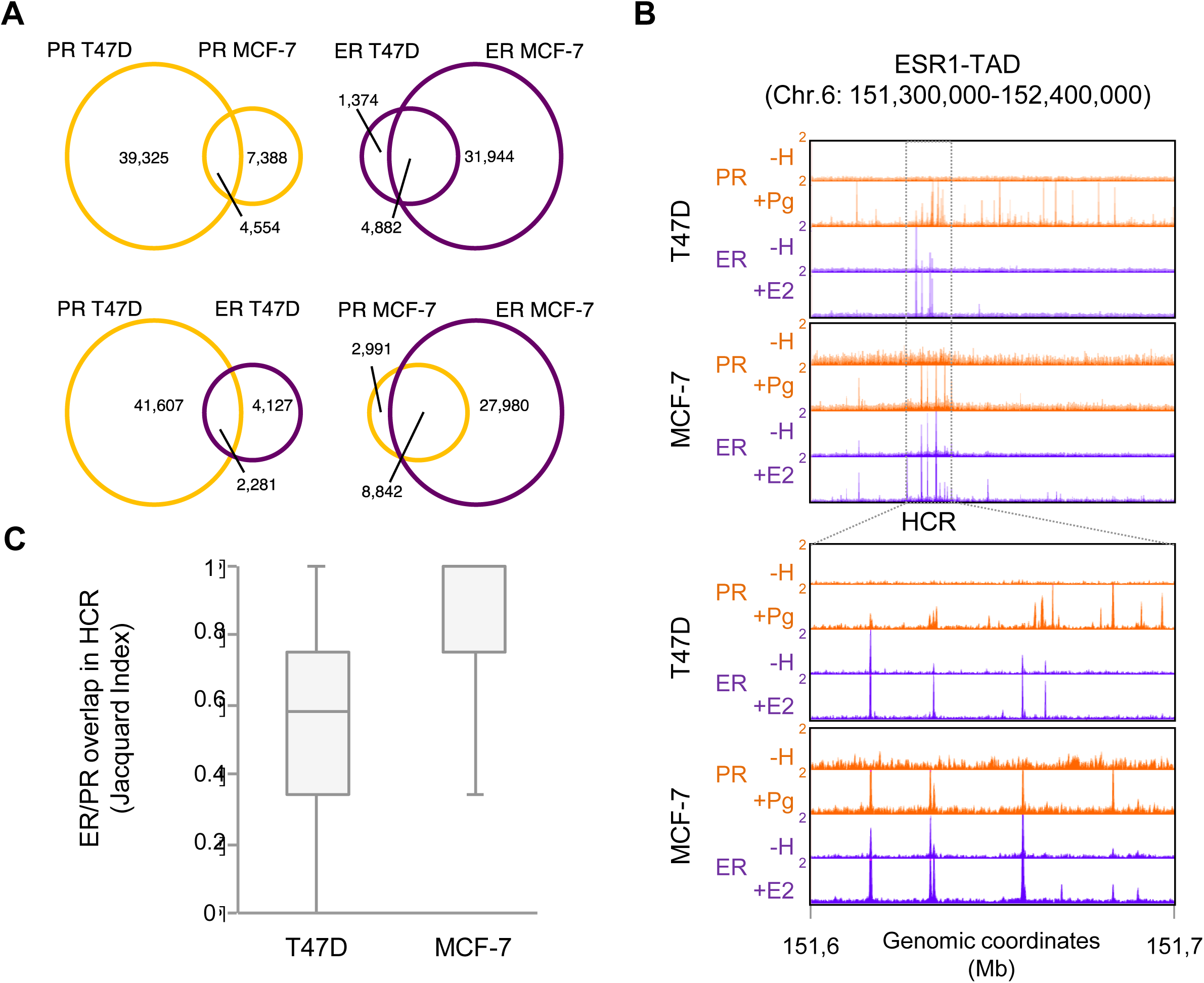
**(A)** Venn diagrams showing the number of PR (top left panel) and ER (top right panel) binding sites overlapping in T47D and MCF-7 exposed to 60 minutes of Pg or E2, respectively. The bottom panels show the overlap between ER and PR binding sites in T47D (left) or MCF-7 (right). **(B)** Genome browser view of ChIP-Seq profiles of PR and ER in T47D (same as in Figure 1) and MCF-7 exposed or not to Progestin (+Pg) or E2 (+E2) for 60 minutes over the ESR1-TAD (top panel). The panel below corresponds to a magnification within the region defined as the ESR1 -TAD HCR for T47D (as in Supplementary Figure 1C) or MCF-7. **(C)** To estimate the overlapping between ER and PR, the Jaccard index between the two types of binding sites was computed for each HCR identified. The boxplot shows the distribution of this index (1 = maximum overlapping) observed in T47D and MCF-7.

**Supplementary figure 3 (related to Figure 3):**
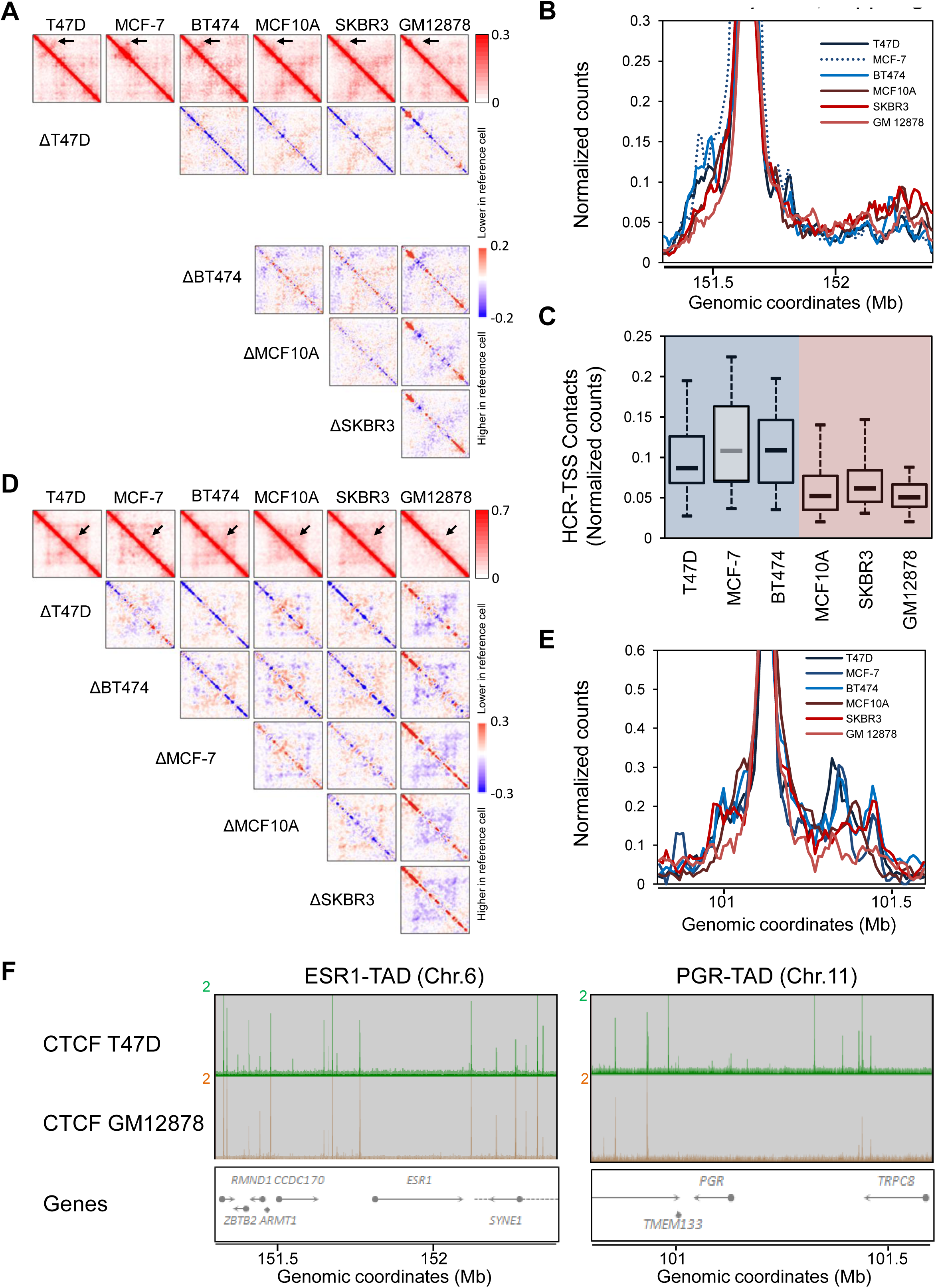
**(A)** Normalised *In Situ* Hi-C contact matrices at 10 kb resolution of the ESR1-TAD (Chr.6: 151,300,000-152,400,000) obtained in the different cell lines grown in the absence of hormones. The matrices below show the differences between the contact matrices of the cell lines at the top minus the reference cell line on the left (blue and red corresponding to contacts higher and lower in the reference cell line, respectively). **(B)** Virtual 4C profiles at 10 kb resolution using the ESR1-TAD HCR as bait in the different ER^+^-PR^+^ (blue lines) and ER^−^-PR^−^ (red lines) grown in absence of hormones. MCF-7 cells (dashed blue line) exhibit a duplication of the HCR region. **(C)** Distribution of contacts between genomic bins within the HCR with the TSS of protein-coding genes within the ESR1-TAD. **(D)** Normalised *In Situ* Hi-C contact matrices at 10 kb resolution of the PGR-TAD (Chr.11: 101,800,000-101,600,000) obtained in the cell lines listed grown in absence of hormones. The matrices below show the differences between the contact matrices of the cell lines at the top minus the reference cell line on the left (blue and red corresponding to contacts higher and lower in the reference cell line, respectively). **(E)** Virtual 4C profiles at 10 kb resolution (expressed as normalized counts per thousand within the region depicted) using the TSS of *PGR* as bait in the different ER^+^-PR^+^ (blue lines) and ER^−^-PR^−^ (red lines) grown in the absence of hormones. **(F)** Genome browser view of ChIP-Seq profiles of CTCF in T47D and GM12878 cells in the ESR1-TAD (left) and PGR-TAD (right).

**Supplementary figure 4 (related to Figure 4):**
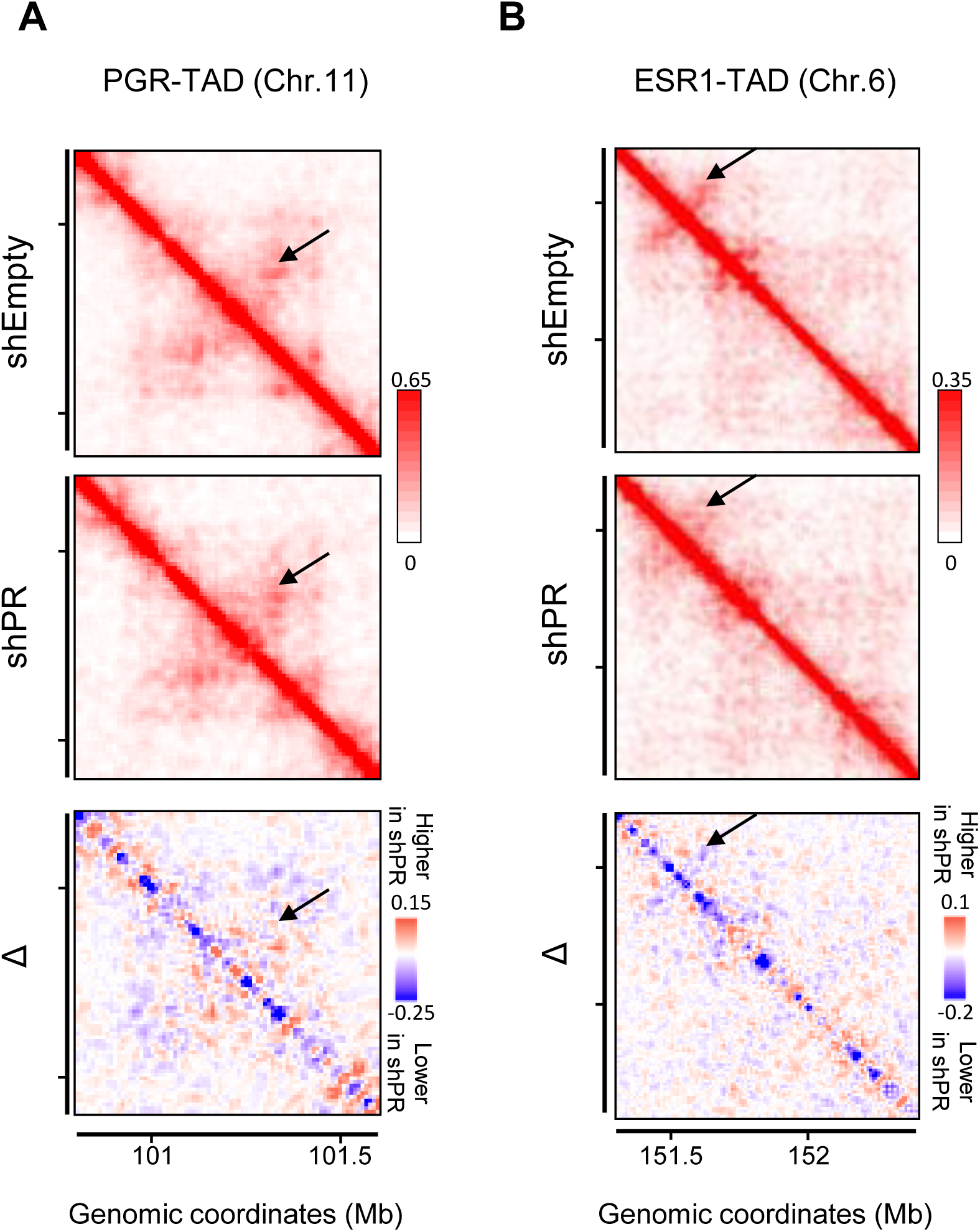
**(A, B)** Normalised *In Situ* Hi-C contact matrices at 10 kb resolution of the region over the ESR1-TAD **(A)** and PGR-TAD **(B)** obtained in T47D expressing control shRNA (shEmpty) or shRNA against PR (shPR). The difference between the contact matrices is shown at the bottom with blue and red corresponding to contacts higher and lower in T47D shEmpty than in T47D shPR, respectively.

**Supplementary figure 5 (related to Figure 5):**
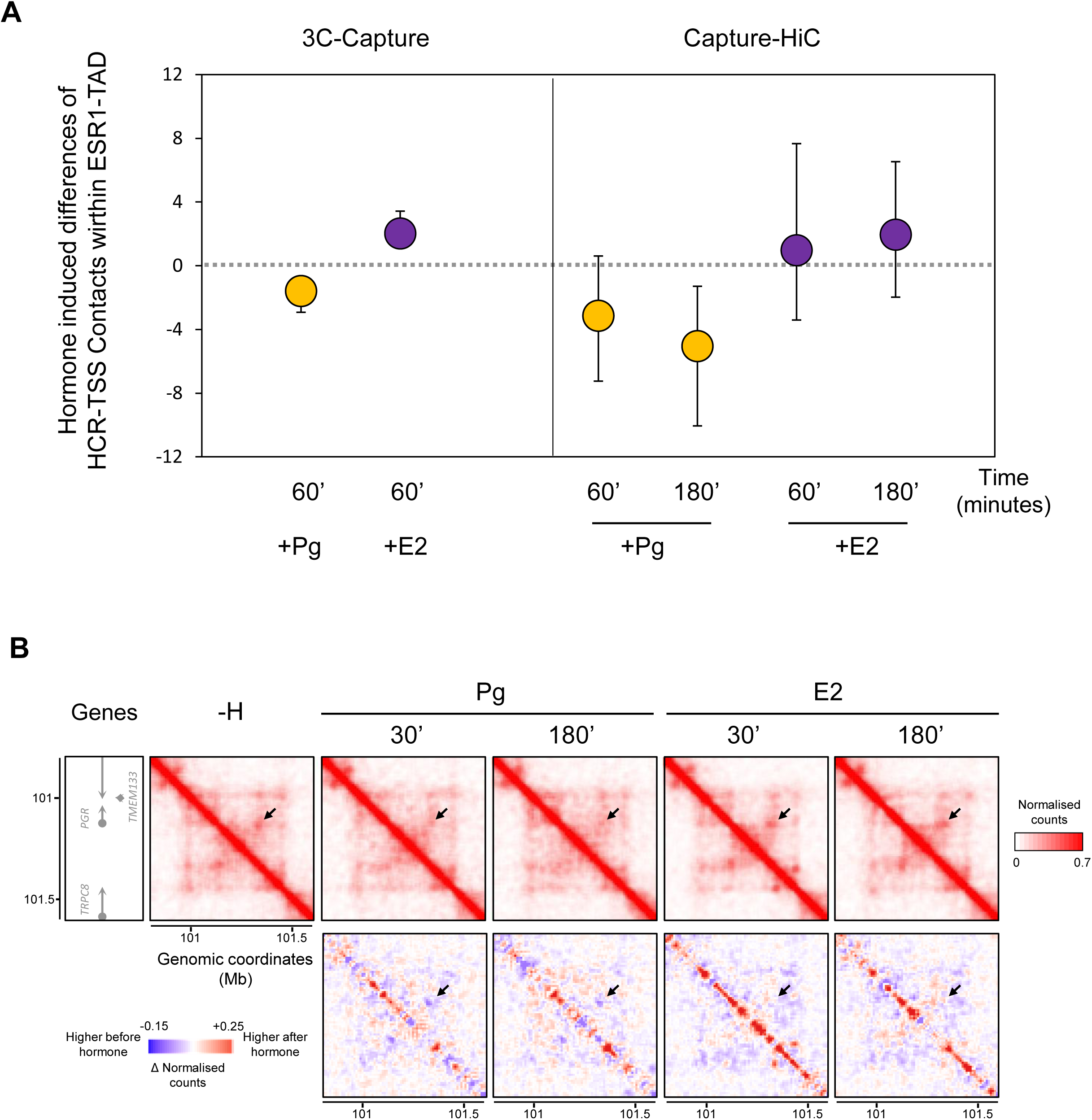
**(A)** Distribution of hormone induced differences of contacts between genomic bins of the HCR with the TSS of protein-coding genes in ESR1-TAD after treatment of T47D cells with Pg (orange) or E2 (purple) in 3C- and Hi-C capture experiments. **(B)** Normalised *In Situ* Hi-C contact matrices at 10 kb resolution of the PGR-TAD obtained in T47D grown in absence of hormone (−H) or after exposure to 30 and 180 minutes to Pg or E2. The matrices below show the differences between the contact matrices compared to the one obtained in absence of hormone (blue and red corresponding to contacts higher and lower in the absence of hormone, respectively).

